# MR-Base: a platform for systematic causal inference across the phenome using billions of genetic associations

**DOI:** 10.1101/078972

**Authors:** Gibran Hemani, Jie Zheng, Kaitlin H Wade, Charles Laurin, Benjamin Elsworth, Stephen Burgess, Jack Bowden, Ryan Langdon, Vanessa Tan, James Yarmolinsky, Hashem A. Shihab, Nicholas Timpson, David M Evans, Caroline Relton, Richard M Martin, George Davey Smith, Tom R Gaunt, Philip C Haycock

## Abstract

Published genetic associations can be used to infer causal relationships between phenotypes, bypassing the need for individual-level genotype or phenotype data. We have curated complete summary data from 1094 genome-wide association studies (GWAS) on diseases and other complex traits into a centralised database, and developed an analytical platform that uses these data to perform Mendelian randomization (MR) tests and sensitivity analyses (MR-Base, http://www.mrbase.org). Combined with curated data of published GWAS hits for phenomic measures, the MR-Base platform enables millions of potential causal relationships to be evaluated. We use the platform to predict the impact of lipid lowering on human health. While our analysis provides evidence that reducing LDL-cholesterol, lipoprotein(a) or triglyceride levels reduce coronary disease risk, it also suggests causal effects on a number of other non-vascular outcomes, indicating potential for adverse-effects or drug repositioning of lipid-lowering therapies.

## Introduction

The continuing success of large scale genome wide associations studies (GWAS) in identifying robust genetic associations,^1, 2^, combined with the development of techniques in Mendelian randomisation (MR)^3, 4^, has vastly expanded the scope for assessing causal relationships between phenotypes^5^. In particular, the exploitation of non-disclosive summary data from GWAS by novel MR methods has been transformative, because this unlocks the powerful feature that causal relationships can be assessed between phenotypes even if these GWASs were performed on non-overlapping samples in a strategy known as two-sample MR^6^ (see Box 1 for further details on MR and its assumptions). The consequence of this methodological development is that the set of potential causal relationships between pairs of phenotypes that can be evaluated grows exponentially, limited only by the availability of reliable GWAS summary data for those phenotypes (Figure 1a).

**Figure 1:**
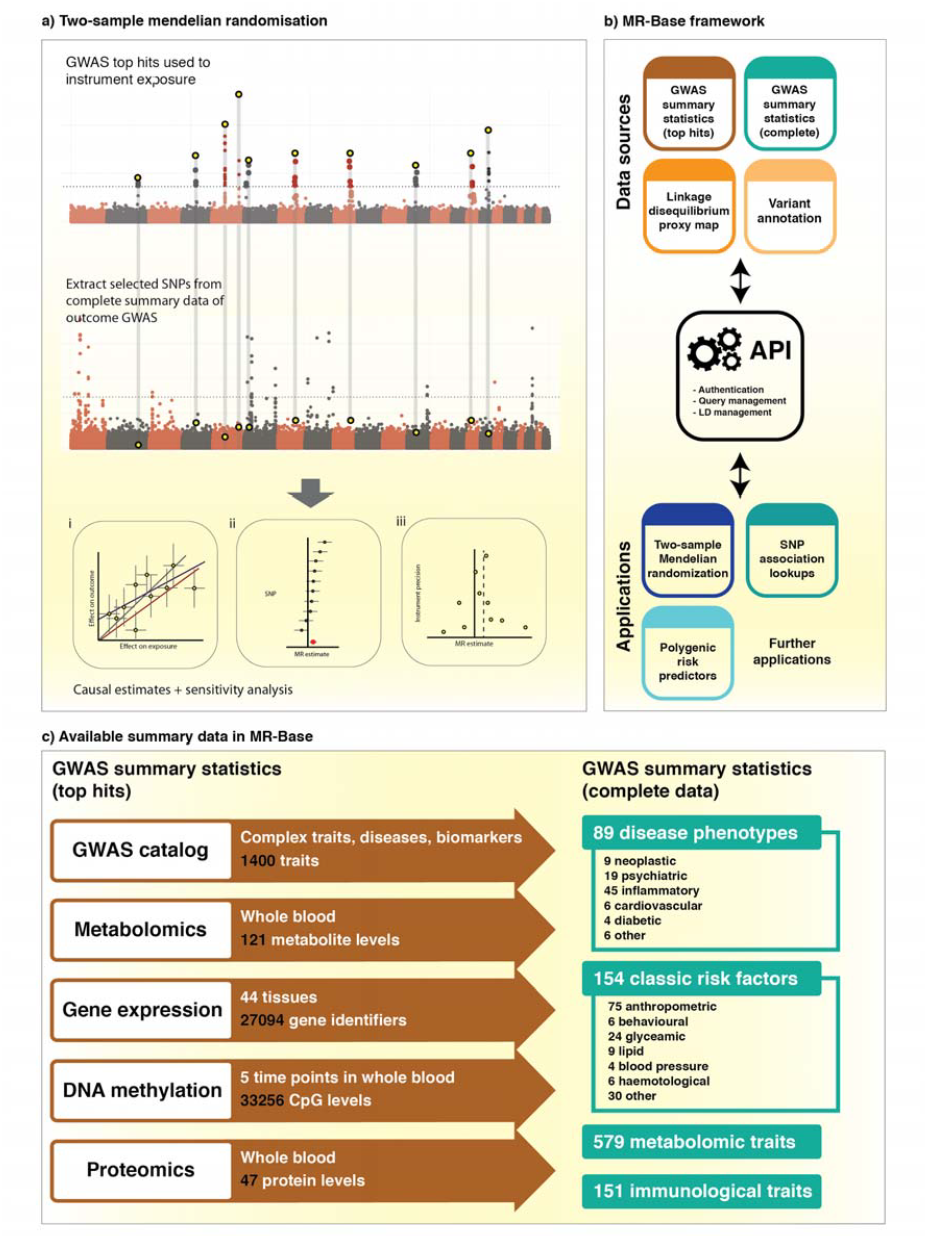
Overview of MR-Base. a) The Two-sample mendel¡an randomisation (2SMR) process for estimating the causal influence of one trait (exposure) on another (outcome) is depicted. Independent SNPs that robustly associate with the exposure are used as instruments (top Manhattan plot). Those SNPs (or LD proxies) are then extracted from the outcome GWAS (bottom Manhattan plot) These summary data can then be harmonised and used to make causal inference using several 2SMR methods (b.i). A range of sensitivity analyses are also performed by default to improve robustness (b.ii and b.iii). All analyses are described in more detail in Supplementary table ##. b) The MR-Base framework consists of modular sources of GWAS summary data, to which access is managed by a public application programming interface (API). The “GWAS summary data (top hits)” module is a resource of data required for genetic instruments. The “GWAS summary data (complete)” module is typically used for outcomes. Applications can interface with the API to use the summary data for a range of analyses. 2SMR is one such analysis which can be performed through an R package or web application, c) A breakdown of the summary data in MR-Base, with the breadth of possible causal inferences that can be made. Any trait for which known GWAS hits are available can be instrumented as an exposure. Any trait for which there is complete summary data can be used as a potential outcome (Supplementary table 1).

The next major challenge is unifying published GWAS summary data with MR methodology within a single analytic platform to begin rapid and systematic interrogation of potential causal relationships across the phenome (Figure 1b). To address this we have developed MR-Base (http://www.mrbase.org/), that exists conceptually as a two-part framework. First, it is a repository of harmonised published GWAS summary data, which has been aggregated from disparate and heterogeneous sources on traits from across the phenome (Figure 1c). The summary data is harvested in two forms, (a) summary data limited to only the statistically significant associations for a particular trait (top hits)^7–11^, and (b) the full set of all SNPs analysed in the GWAS of a trait (complete summary data). The utility of these different forms of summary data go beyond making causal inference, but their use in MR is described in Figure 1. Second, MR-Base plays host to a range of causal estimation methods and automatically applied sensitivity analyses that can be used to improve the reliability of causal inferences^6, 12–19^. The data and the methodology repositories are curated and continually expanding.

We showcase the application of MR-Base through a systematic MR study of the effect of LDL cholesterol, lipoprotein(a) [Lp(a)] and triglycerides-intervention targets for the prevention of coronary heart disease (CHD)-on a range of health-related outcomes. First, We perform MR analyses to predict the efficacy^20^ and safety^21,22^ of emerging lipid-lowering drugs^23,24^. Second, to gain insights into the broad safety of lipid-targeted interventions, we perform MR analyses of lipids against a wide range of disease outcomes and related traits. These examples demonstrate one particular scope of MR-Base: predicting the downstream consequences of interventions on particular phenotypes.

### Box 1: Mendelian randomization and instrumental variables

The underlying principle of Mendelian randomization (MR) is that if a genetic variant (G) affects an environmental, molecular or physiological exposure (e.g. smoking, low density lipoprotein [LDL] cholesterol levels, body mass index), it can be used as an instrumental variable or proxy to appraise the causal effect of that exposure on an outcome at a population level. In this paper we refer to the trait that is the putative causal factor as the ‘exposure’ (X) and the trait that is consequential (e.g. a disease or other complex trait) as the ‘outcome’ (Y), following conventions in the epidemiological literature.

Genetic variants are good potential instrumental variables because they are fixed from conception and tend to be randomly distributed in the general population with respect to lifestyle and environmental factors. Thus, studies of gene-phenotype associations are less susceptible to confounders (U) and reverse causation in comparison to conventional observational studies. If MR assumptions are met, associations of a genetic instrument with an exposure and outcome can be used to infer the causal effect of the exposure on the outcome. The assumptions are: (IV1) the instrument is associated with the exposure; (IV2) the instrument is independent of known and unknown confounders; and (IV3) the instrument is independent of the outcome conditional on the exposure and confounders (Figure below and supplementary table 10). For further details see ref^4^. Both G-X and G-Y can be estimated using summary association statistics from non-overlapping GWAS studies, in an approach known as two-sample MR^6^.

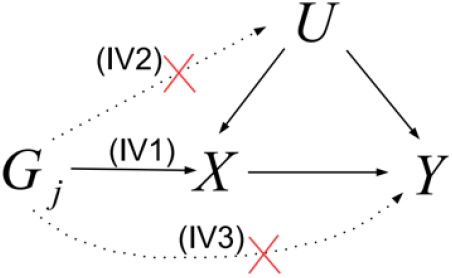

Interpretation of a MR study requires careful consideration of whether genetic variants exhibit horizontal pleiotropy, i.e. they associate with the exposure but influence the outcome through pathways that do not include the exposure. This is a violation of assumption IV3. Using multiple instruments for an exposure, where possible, can help to evaluate and potentially account for this assumption. A range of analyses are now available to assess the sensitivity of results to horizontal pleiotropy, including MR Egger regression^18^ and the weighted median function^19^ (Supplementary figure 8, Supplementary table 2 and Supplementary table 10). Although MR can provide evidence regarding causality, we strongly advocate the use of different study designs in order to triangulate evidence when making causal inferences^25^.

## Results

### The MR-Base resource

We curated and harmonised ‘complete summary data’ from 1094 GWAS analyses, corresponding to 139 unique GWAS publications and 61 unique studies (including 44 consortia) (Figure 1c and Supplementary table 1). At the time of writing (November 2016), these GWAS summary statistics correspond to approximately 4 billion SNP associations with 56 diseases, 125 risk factors, 559 metabolites and 149 immune system traits, derived from GWAS analyses of ∼14.9 million individual sample measurements. In addition to the ‘complete summary data’, we also collected published GWAS associations that comprise only the significant hits of a GWAS after applying stringent p-value thresholds (e.g. 5E-8, a conventional threshold for declaring statistical significance in GWAS) and often performing replication to obtain unbiased effect sizes. These ‘top hits’, which can be used to define genetic instruments for exposures in MR analyses (see Box 1), include 22,369 SNPs associated with 3889 complex traits and diseases in the NHGRI-EBI GWAS catalog^7^; 187,318 SNPs associated with DNA methylation levels in whole blood at 33,256 genomic CpG sites, across five time points^8^; 187,263 SNPs associated with gene expression levels at 27,094 gene identifiers, across 44 different tissues^9^; 1088 SNPs associated with metabolite levels in whole blood for 121 different metabolites^10^; and 56 SNPs associated with protein levels in 47 different analytes^11^ (Online methods).

The platform integrates these summary associations with statistical techniques for MR, including inverse-variance weighted (IVW) linear regression^12,13^ (the recommended default), maximum likelihood^6^ and Wald ratio methods^26^, as well as sensitivity analyses that allow users to assess the potential for violations of MR assumptions (Supplementary table 2), thereby improving the reliability of causal inference. An example of the basic use case of MR-Base-assessing the causal relationship between two traits-is presented in the Supplementary Note, where we reproduce the known^27,28^ causal effect of higher LDL cholesterol levels on elevated risk of CHD.

### Predicting the efficacy and safety of lipid lowering drugs

In order to predict the efficacy of current LDL cholesterol-lowering drugs for cardiovascular disease prevention, we used genetic variants at the 3-Hydroxy-3-Methylglutaryl-CoA Reductase (*HMGCR*), Niemann-Pick C1-Like 1 (*NPC1L1*) and Proprotein convertase subtilisin/kexin type 9 (*PCSK9*) genes to mimic the action of statins, Ezetimibe and Evolocumab, respectively, and compared the results to findings from randomized controlled trials (RCTs)^20,21,28,29^ (Figure 2, Supplementary table 3). Odds ratios (ORs) for CHD per standard deviation (SD) decrease in LDL cholesterol were directionally similar in MR and RCT analyses and were indicative of reductions in disease risk: 0.61 (95% confidence interval [Cl]): 0.48-0.79) vs. 0.76 (0.74-0.77) for HMGCR/statins; 0.65 (0.40-1.06) vs. 0.85 (0.75-0.97) for NPC1L1/Ezetimibe; 0.41 (0.30-0.56) vs. 0.67 (0.51-0.88) for PCSK9/Evolocumab.

**Figure 2.**
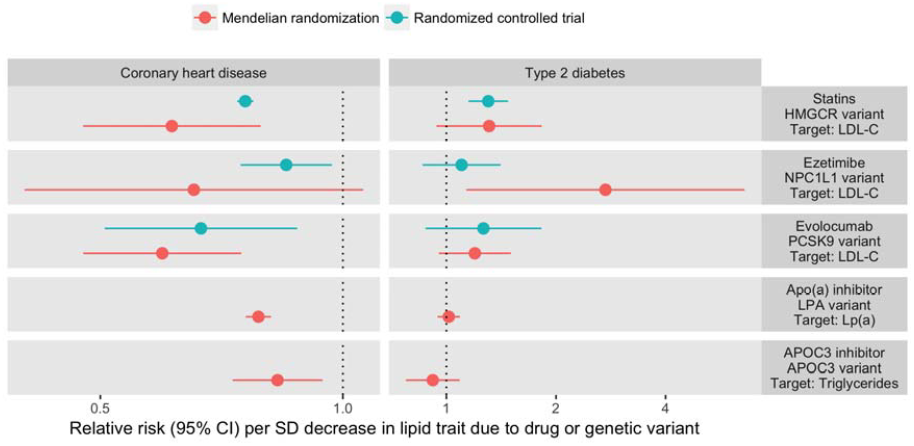
Effect of lipid lowering on risk of coronary heart disease and type 2 diabetes in randomized controlled trials or predicted by Mendelian randomization. Each point represents the relative risk for disease per each standard deviation increase in selected lipid traits due to the intervention (green) or genetic pathway (red). Results are not yet available for drug trials of apo(a) and AP0C3 inhibitors. Hazard ratios (from clinical trial results) and odds ratios (from Mendelian randomization analyses) were assumed to approximate the same measure of relative risk. The number of disease events from clinical trials of statins were 24323 CHD^28^ & 7339 type 2 diabetes^20^ cases; from trials of Ezetimibe were 5314 CHD^29^ & 1414 Type 2 diabetes^29^ cases; from trials of Evolocumab were 60 CHD^21^ & 45 type 2 diabetes^21^ cases. The number of cases in Mendelian randomization analyses of CHD were 60801^63^ and for type 2 diabetes were 26488^64^. Inhibitors of HMGCR, NPC1L1 and PCSK9 reduce LDL cholesterol concentration; inhibitors of apo(a) reduce Lp(a) concentration; inhibitors of AP0C3 reduce triglyceride concentration. Effects of SNPs on LDL cholesterol and triglycerides were estimated using results from the GLGC^6l^; The effect of rs10455872 on Lp(a) levels was estimated using results from the European Prospective Investigation into Cancer and Nutrition (EPIC) study^60^. **Abbreviations**: **CHD**, coronary heart disease; **SD**, standard deviation; **Cl**, confidence interval; **LDL**, lipoprotein; **Lp(a)**, lipoprotein(a); **Apo(a)**, apolipoprotein(a); ***HMGCR***, 3-Hydroxy-3-Methylglutaryl-CoA Reductase gene; ***NPC1L1***, Niemann-Pick C1-Like 1 gene; ***PCSK9***, Proprotein convertase subtilisin/kexin type 9 gene; ***LPA*** Lipoprotein (A) gene; ***APOC3***, Apolipoprotein C3 gene.

We used the same framework to predict the efficacy of two novel lipid-lowering drugs that are currently undergoing clinical trials: an oligonucleotide inhibitor of apolipoprotein(a) *(APOA)* that targets Lipoprotein(a) (Lp[a]), and an oligonucleotide inhibitor of apolipoprotein C3 *(APOC3)* that targets triglyceride levels. In MR analyses there was strong evidence that intervention at both novel targets is likely to reduce risk of CHD. The OR for CHD per SD decrease in Lp(a) due to APOA was 0.79 (0.76-0.81) and per SD decrease in triglycerides due to APOC3 was 0.81 (0.73-0.90) (comparable results from RCT studies were unavailable).

To evaluate the safety of these lipid-lowering drugs, we went on to estimate associations with type 2 diabetes risk, a well known side-effect of statin treatment (Figure 2, Supplementary table 3). ORs for type 2 diabetes per SD decrease in LDL cholesterol were directionally similar in MR and RCT analyses: 1.31 (0.94-1.82) vs. 1.30 (1.15-1.50) for HMGCR/statins; 2.73 (1.13-6.59) vs. 1.10 (0.86-1.41) for NPCIL1/Ezetimibe; 1.20 (0.95-1.50) vs. 1.26 (0.88-1.82) for PCSK9/Evolocumab. In MR analyses, the OR for type 2 diabetes per SD lower Lp(a) due to APOA was 1.02 (0.95-1.09) and per SD lower triglycerides due to APOC3 was 0.88 (0.76-1.02) (comparable results from RCT studies were not available).

LDL-cholesterol targeting drugs could increase type 2 diabetes risk through LDL-cholesterol, or through alternative pathways (horizontal pleiotropy) specific to the target loci. To test this, we performed MR analyses of LDL-cholesterol using all available instruments (Supplementary table 3) against type 2 diabetes (Figure 3, Supplementary table 4). The IVW method estimated a strong influence of higher LDL-cholesterol levels on type 2 diabetes risk (1.23 [1.11-1.36] OR per SD increase), with similar estimates obtained from MR Egger and weighted median analyses (1.39 (1.17-1.65) and 1.31 (1.16-1.48), respectively). These results show that lower circulating LDL-cholesterol may increase risk of type 2 diabetes, suggesting that intervention on any gene that influences LDL-cholesterol levels is liable to incur this increased risk.

**Figure 3:**
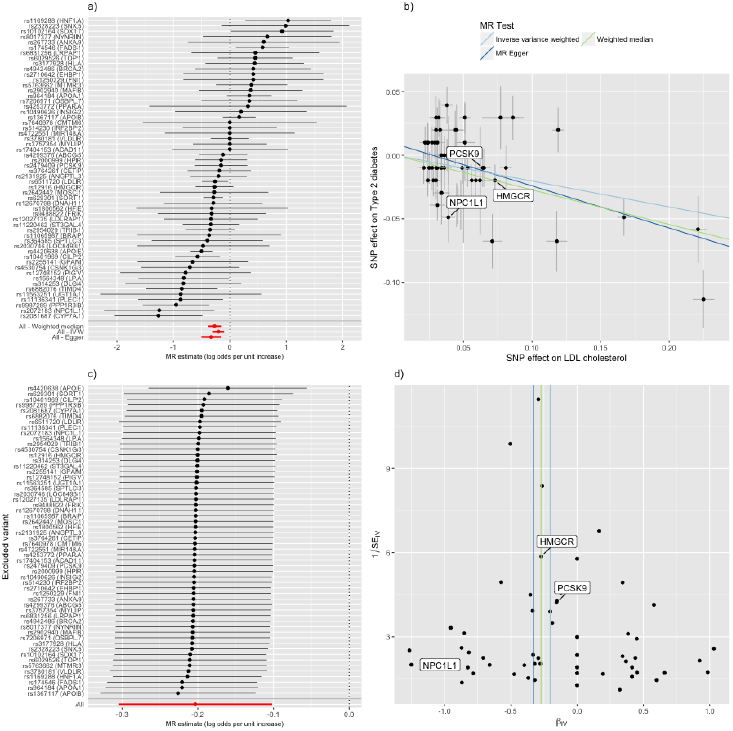
Detailed analysis of causal effect of LDL cholesterol (LDL-C) on type 2 diabetes (T2D). a) A forest plot, where each black point represents the causal estimate of LDL-C (SD units) on T2D {log(OR)) produced using each of the 57 instruments separately, and red points show the combined causal estimate using all SNPs together using each of three different methods. Horizontal lines denote 95% confidence intervals, b) A plot relating the effect sizes of the SNP-LDL association (x-axis, SD units) and the SNP-T2D associations (y-axis, log{OR)) with 95% confidence intervals. The slopes of the lines correspond to causal estimates using each of three different methods, c) Leave-one-out sensitivity analysis. Each black point represents the maximum likelihood MR method applied to estimate the causal effect of LDL-C on T2D excluding that particular variant from the analysis. The red point depicts the estimate using all SNPs. All estimates in this plot are obtained using the inverse variance weighted method. There are no instances where the exclusion of one particular SNP leads to dramatic changes in the overall result, d) Funnel plot showing the relationship between the causal effect of exposure on outcome estimated by each SNP against the inverse of the standard error of the causal estimate. Vertical lines show the MR estimates using all SNPs for each of four different methods. Symmetry of the funnel plot indicates lower risk of horizontal pleiotropy leading to unreliable associations.

### Predicting the impact of lowering lipid levels on human health

In order to gain broader insight into the effect of lowering lipid levels on health outcomes more generally we conducted a hypothesis-free scan for causal influences of LDL cholesterol (instrumented using 57 SNPs), Lp(a) (instrumented using 1 SNP) and triglycerides (instrumented using 40 SNPs) on 40 non-vascular diseases and 108 non-lipid complex traits in MR-Base (Figure 4, Supplementary figure 2). In addition, using an unadjusted p-value of 0.05 to denote suggestive evidence for association, we identified 16 and 17 out of 147 non-vascular traits associated with LDL cholesterol and triglycerides, respectively, and 11 out of 112 non-vascular traits for Lp(a). We went on to examine the reliability of these 33 putative associations for LDL cholesterol and triglycerides in greater detail using a range of sensitivity analyses (Supplementary figure 3).

**Figure 4.**
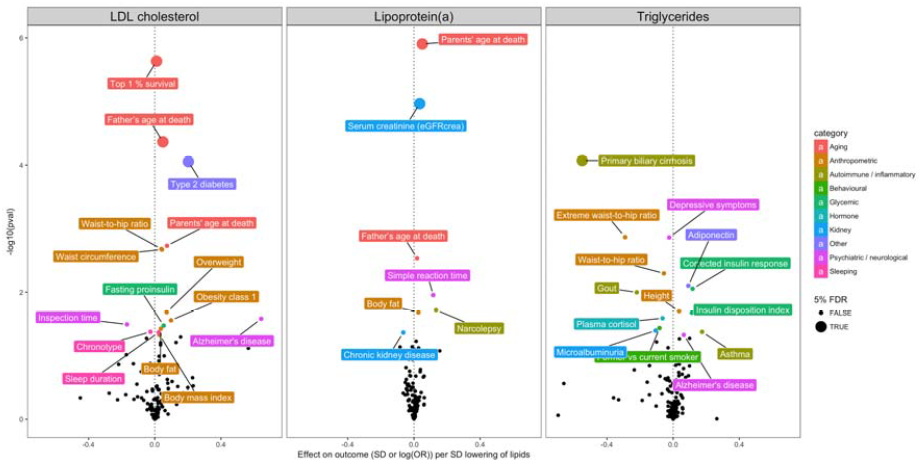
Hypothesis-free causality scan, a) An illustration of the analysis that was performed. LDL cholesterol, lipoprotein(a) and triglycerides were instrumented by 57, 1 and 40 SNPs, respectively. MR was performed sequentially for each exposure on each of 150 complex trait outcomes. The word cloud lists the categories under which each complex trait falls, with text size corresponding to the number of traits for that category, b) Volcano plot showing the effect of lower lipid levels on outcomes (x-axis) against the-logl0(p-value) (y-axis) obtained from the inverse variance weighted estimate. Each volcano plot represents the association of a particular exposure against all available outcomes. Those outcomes that have a p-value < 0.05 are labelled. Larger points denote false discovery rate (FDR) < 0.05.

#### Associations surviving multiple testing correction

MR associations surviving multiple testing correction (false discovery rate, < 0.05) for LDL cholesterol were type 2 diabetes (described earlier), and two traits for which lower LDL levels related to increased longevity: top 1% survival and father’s age at death. For these outcomes the MR estimates were consistent using IVW, MR Egger and weighted median analysis providing strong evidence that reducing LDL cholesterol levels increases longevity.

Lower triglycerides appeared to have a strong association with reduced risk of primary biliary cirrhosis (PBC). The MR estimates using IVW, MR Egger and weighted median analysis were consistent, though the effect slightly attenuated upon exclusion of the *GCKR* genetic variant (a gene with known pleiotropic effects) in leave-one-out analyses.

Lower Lp(a) associated strongly with parents’ age at death, similar to LDL-C. Kidney function also appeared to be influenced, with lower Lp(a) associated with higher serum creatinine levels. Consistent with this finding, risk of chronic kidney disease was reduced by lower Lp(a). Because Lp(a) was only instrumented by a single SNP further sensitivity analyses were not possible.

#### Putative associations not surviving multiple testing correction

The apparent association between lower LDL cholesterol and increased risk of Alzheimer’s disease using the IVW method was not reliable. MR Egger and weighted median methods attenuated strongly to the null, and after exclusion of the *APOE* genetic variant (rs4420638) in leave-one-out sensitivity analyses the IVW estimate attenuated to the null also. The putative association of lower triglycerides on Alzheimer’s disease, however, remained consistent between the three different MR methods, though precision reduced substantially upon removal of the *VEGFA* genetic variant (rs998584) in leave-one-out sensitivity analyses.

The weak putative association between triglycerides and asthma was not consistent amongst the IVW, MR Egger and weighted median methods. The asymmetry of the funnel plot suggested some level of heterogeneity, but there were only 18 out of 40 variants available (after searching for LD proxies) in the asthma summary data which likely contributes to reduced statistical power.

Putative associations also appeared for lower LDL cholesterol with higher anthropometric measures (body mass index, risk of obesity, body fat, waist-to-hip ratio and waist circumference). In all cases the MR Egger and weighted median estimates were stronger than the IVW estimates. However, in leave-one-out analyses for each of these MR analyses the effects attenuated towards the null upon exclusion of the *APOE* genetic variant.

## Discussion

As the availability of published GWAS summary data continues to grow MR offers an increasingly attractive approach for exploring the aetiology of disease. We have developed a platform that integrates a database of GWAS summary data together with a public repository of statistical methods for enabling systematic causal inference across the phenome. This benefits modelling of phenomic relationships in two ways-first, it maximises the breadth of possible causal relationships that can be interrogated by drawing together genetic information on as many traits as possible. Second, automating the application of state-of-the-art methodology establishes core standards for reporting MR results and improves the reliability and reproducibility of causal inference.

One important emerging area in MR is predicting the safety and efficacy of drug targets for for disease prevention^30^. We recapitulate the known beneficial effects of LDL-cholesterol lowering on coronary disease risk^31,32^, and the small risk-raising effect on type 2 diabetes^20,22,23^. Although our study was limited to publicly available data for the analyses of PCSK9, NPC1L1 and HMGCR and type 2 diabetes, our results are similar to, though less precise than, findings from recent larger MR studies^22,34,35^. The latter used the same public datasets that support MR-Base combined with additional data and stronger instrumental variables, illustrating how future studies could combine the convenience of MR-Base with novel datasets to maximise statistical power.

The associations of LDL-cholesterol lowering with reduced CHD risk and increased type 2 diabetes risk were observed regardless of whether we considered specific gene targets (HMGCR, PCSK9 and NPC1L1) separately or combined all LDL lowering genetic variants together in a single instrument. These findings suggest that the effect of LDL-lowering drugs on cardiometabolic disease is, at least partly, due to an on-target mechanism and that interventions through any gene that influences LDL-cholesterol levels is likely to increase risk of type 2 diabetes and decrease risk of CHD. The full mechanism of how LDL-targeted interventions raise risk of type 2 diabetes remains to be elucidated, but our hypothesis-free scan highlighted the potential role for anthropometric markers, such as body mass index.

This is consistent with evidence from RCT and MR studies indicating that statin treatment is associated with weight gain, a well known risk factor for type 2 diabetes^20,33^.

We also use MR-Base to predict the impact of oligonucleotide-inhibitors of APOC3 and APOA: effective pharmaceutical strategies for lowering triglycerides^23^ and Lp(a)^36,37^ levels, respectively, but with, as yet, unknown efficacy for preventing coronary disease and unknown safety profile. Our findings indicate that these novel pharmaceutical agents are likely to be effective at reducing risk of CHD, with little evidence for adverse effects on risk of type 2 diabetes. However, these results cannot address the potential for adverse effects due to off-target mechanisms.

In a hypothesis-free screen, we found evidence for effects of lipid-lowering on non-cardiometabolic outcomes. Lower Lp(a) concentration was associated with higher serum creatinine levels. Lower serum creatinine is a diagnostic factor for chronic kidney disease, which also showed some evidence for association with higher Lp(a) levels in our analyses, consistent with observational studies^38–40^. These results suggest some potential for the repositioning of APOA-inhibitors for the prevention of chronic kidney disease.

The association between triglycerides and primary biliary cirrhosis (PBC) is well known observationally^41^ but the association is assumed to be the consequence of the disease. Our results suggest, however, that elevated triglyceride concentration is also a contributing causal factor for primary biliary cirrhosis. To test if the reverse causal effect (PBC influencing triglycerides) also exists, we performed a further MR analysis using 25 variants^42^ to instrument PBC against triglycerides, which gave a weak positive association (Supplementary figure 4), which though low in precision is in the direction expected based on observational associations. Further studies are required to confirm this result and to explore whether triglyceride lowering could be a useful treatment strategy for disease prevention. One concern, which should be considered in any MR application, is the possibility of spurious associations arising due to selection bias^43^. Here for example, the triglyceride-PBC association may have arisen if individuals with PBC had been excluded from the triglycerides GWAS. Further exploration suggests that this mechanism is unlikely in this specific case because PBC is sufficiently rare to not have a large impact on the triglycerides GWAS.

LDL cholesterol and Lp(a) were also associated with parents’ age at death. These results potentially reflect the influence of these lipid traits on CHD risk, given that CHD is the single leading cause of death in the world^44^.

A consequence of this scale of automation is that the problem of causal inference shifts from handling complex data to interpreting complex results. In our hypothesis-free scan for downstream effects of higher lipid levels, though we perform many tests we do still find associations that survive correction for multiple testing. GWAS sample sizes continue to grow and power to detect any one particular association is improving. But as the scale of phenomic data mining grows to include more traits, multiple testing will make the traditional approach of identifying associations increasingly intractable. Developing modelling frameworks that enable the construction of large causal systems comprising small putative effects is likely to be warranted.

GWAS is typically conducted using meta-analysis of large numbers of different cohorts, where each cohort is likely to contribute to GWAS on many different traits. As a consequence there is likely to be considerable sample overlap amongst different GWAS within the database. In two-sample MR non-independence of samples for the exposure and outcome GWAS can induce bias of causal estimates towards the confounded observational association if instruments are weak (canonically defined as having SNP-exposure associations with F-statistics < 10 but the bias various continuously with instrument strength). If samples are independent, however, weak instruments induce bias towards the null. The level of sample overlap amongst different studies is not yet quantified within the database, but in addition to manual curation this could also be inferred analytically using LD score regression.

There exist several resources for the collection and curation of genetic summary data. The NHGRI-EBI GWAS Catalog^7^ prioritises published associations that are statistically significant, according to conventional GWAS thresholds, and show evidence for replication in independent datasets. GWASdb^45^ and GRASP^46^ similarly focus on published association statistics but employ more relaxed P-value inclusion criteria. We use the GWAS catalog in MR-Base as a source of instruments for potential exposure variables. Other important resources include PhenoScanner^47^, which collates complete summary data from 637 GWAS datasets, and dbGAP, which collates individual and complete summary-level genetic data (but in a generally less accessible and less standardized format). MR-Base has several notable differences to these existing repositories of GWAS summary data: comprising 1094 datasets (as of December 2016), it is larger than PhenoScanner, more easily accessible than dbGAP, does not employ P-value inclusion thresholds (unlike the GWAS catalog, GWASdb and GRASP), provides a security layer which enables data to be deposited with restricted access (e.g. for unpublished studies) and it integrates directly to software to automate implementation of causal inference through two-sample MR. MR-Base also actively seeks datasets from studies that have yet to release their full results into the public domain (these currently correspond to 4% of the 1094 datasets in MR-Base). A number of other resources are now available that integrate published data of various (often non-GWAS) experimental origins with analytical methods, especially for the identification of potential drug targets, for example Open Targets^48^ and DisGeNET^49^ to name but a few. Combining these different resources with results from MR could be a valuable approach towards triangulating evidence about causal relationships in order to improve reliability of findings^25^.

The data behind MR-Base can easily be extended to accommodate other MR methods, such as network MR^51^, or other other post-GWAS analytical approaches such as fine mapping^52–54^, constructing genetic predictors^55^, omic-wide association studies^2^, or understanding genetic architecture^56,57^. For example, the GWAS summary statistics used in MR-Base also support LD Hub^58^, an online application for calculating bivariate genetic correlations and trait heritability using LD score regression (http://ldsc.broadinstitute.org/).

In conclusion, we have developed MR-Base: a framework for a) collating and harmonising GWAS summary associations statistics and b) harnessing GWAS summary association statistics to automate implementation of two-sample MR. MR-Base is a growing, collaborative and open source platform that maximises the potential of summary-level genetic data for causal inference across the phenome.

## Online methods

### Overview

MR-Base comprises two main components: a database of GWAS summary association statistics and LD proxy information, and an R package that serves as repository for MR methods and sensitivity analyses (TwoSampleMR, https://github.eom/MRCIEU/TwoSampleMR). The database is accessible through an application programming interface (API), which enables optional access restrictions for summary data that is not released publicly. A web app was developed as a user-friendly wrapper to the R package using the R/shiny framework. Scripts to perform the analyses presented in this paper are available at https://github.com/explodecomputer/mr-base-methods-paper.

### MR-Base database

#### Obtaining summary data from genome wide association studies

We downloaded publicly available datasets from study-specific websites and dbGAP and invited studies curated by the GWAS catalog to share data (if these were not already publically available) (Supplementary figure 4). To be eligible for inclusion in MR-Base, studies must provide the following information for each SNP: the beta coefficient and standard error from an additive regression model and the modelled effect and non-effect alleles. This is the minimum information required for implementation of two-sample MR. The following information is also sought but is not essential: effect allele frequency, sample size, P values for SNP-phenotype associations, P values for Hardy–Weinberg equilibrium, P values for Cochran’s Qtest for between study heterogeneity (if a GWAS meta-analysis) and metrics of imputation quality, such as info or r^2^ scores (for imputed SNPs). MR-Base also includes information on the following study-level characteristics: sample size, number of cases and controls (if a case-control study), standard deviation of the sample mean for continuously distributed traits, geographic origin and whether the GWAS was conducted in males or females (or both). In future extensions to MR-Base, we plan to collate more detailed information on phenotype distribution (e.g. sample means for continuously distributed phenotypes) and population characteristics (mean and standard deviation of age, number of males and females) and statistical models (e.g. covariates included in regression models and genomic control inflation factors).

#### Linkage disequilibrium proxy data

One of the main functions of the MR-Base database is to extract association data for requested SNPs from GWAS studies of interest to the user (Association Table, Supplementary figure 7). Often, however, a requested SNP may not be present in the requested GWAS (e.g. because of different imputation panels or because imputed SNPs were not available). In order to enable information to be obtained even when SNPs are missing, we provide an LD proxy function using 1000 genomes data from 503 European samples. For each common variant (minor allele frequency [MAF] > 0.01) we used plink1.90 beta 3 software to identify a list of LD proxies. We recorded the r^2^ values for each LD proxy, the phase of the alleles of the target and proxy SNPs. We limited the LD proxies to be within 250kb or 1000 SNPs and with a minimum r^2^=0.6.

### MR instrument catalog

We have assembled a collection of potential instruments for a wide range of traits from various sources. These sources typically present only the top hits from a GWAS, rather than the entire GWAS summary statistics. As such, the traits included here only have sufficient data for them to be evaluated as potential exposures. All curated data is available through the MRInstruments R package (https://github.com/MRCIEU/MRInstruments):

### NHGRI-EBI GWAS catalog

This is a comprehensive catalog of reported associations from published GWAS studies^7^. To harmonize the data to be suitable for MR we converted odds ratios into log odds ratios, inferring standard errors from reported 95% confidence intervals or (if the latter were unavailable) from the reported P value using the Z distribution. We extracted information on the units of the SNP-trait effect; identifying the effect and non-effect alleles, by comparing the risk allele reported in the GWAS catalog to allele information downloaded from ENSEMBL, using the R/biomaRt package^59^. R/biomaRt was also used to identify base pair positions (in GRCh38 format) and associated genes; and infer effect allele frequency from the risk allele frequency reported in the GWAS catalog. We excluded SNP-trait associations from the GWAS catalog if they were missing a P value, beta (estimate of the SNP-trait effect) or a standard error for the beta. The MR-Base standardized version of the GWAS catalog (2016-07-17 release at time of writing) comprises 24,305 potential instruments for approximately 4960 traits. The potential of this catalog for MR is considerable, but there are issues. For example not all reported associations surpass a P value threshold of 5e-8; and effect sizes are not always presented in the same units within studies of the same trait, hence further curation is warranted. MR results generated using the GWAS catalog should be viewed with caution.

### Accessible Resource for Integrated Epigenomics Studies (ARIES) mQTL catalog

We obtained a large set of instruments for DNA methylation levels that were estimated in the ARIES dataset ^8^. Methylation quantitative trait loci (mQTLs) were identified in 1000 mothers at two time points and 1000 children at three time points. Top hits were obtained from http://mgtldb.org with P<1e-7. There are 33,256 unique CpG sites across the 5 time points with at least one independent instrument.

### GTEx eQTL catalog

We used the GTEx resource ^9^ of published independent cis-acting expression QTLs (cis-eQTLs) to create a catalog of SNPs influencing up to 27,094 unique gene identifiers across 44 tissues.

### Metabolomic QTL catalog

SNPs influencing 121 metabolites measured using nuclear magnetic resonance (NMR) analysis in whole blood were obtained ^10^, totalling 1088 independent QTLs across all metabolites.

### Proteomic QTL catalog

SNPs influencing 47 protein analyte levels^11^ in whole blood were obtained, totaling 57 independent proteomic QTLs.

The above catalogs can be used to define the user’s instruments in an MR analysis. Alternatively, the user can define their instruments manually (e.g. by uploading a file to the website) or can use the MR-Base repository of full GWAS summary association statistics to extract independent sets of variants that surpass user-specified P-value and clumping thresholds.

## Applied MR analyses

Instruments for LDL cholesterol levels and triglyceride levels were obtained from reported independent significant associations by the Global Lipids Genetics Consortium (GLGC) GWAS (n = 173,044), resulting in 57 and 40 independent instruments, respectively (Supplementary table 3). We used rs10455872 as an instrument for natural log Lp(a) levels, based on findings in a previous study^60^. These rs IDs were extracted from the complete summary data of 42 diseases and 108 non-lipid risk factors. Where the exact rs ID wasn’t available an LD proxy was searched for with r^2^> 0.8 and within 250kb of the target SNP. The effect alleles of each SNP used in the exposure and outcome GWAS were compared and converted to the same strand where discrepancies were observed. Effect sizes in the exposure and outcome GWAS had their directions switched to reflect the same effect allele. All effect sizes were in standard deviation units for continuous traits or log odds ratios for disease and binary traits. We primarily used the two-sample MR IVW method to report causal estimates between exposures and outcomes. Where indicated, we also used MR-Egger regression and the weighted median function to assess the sensitivity of our results to violations of MR assumptions. The diseases and complex traits specified as outcomes in the MR analyses are described in Supplementary table 5.

To predict the effects of lipid lowering drugs on health outcomes, we used SNPs with known associations with LDL cholesterol, triglyceride and Lp(a) levels that were within the vicinity of the genes that are targeted by specific drugs. Statin drug effects, targeting the *HMGCR* gene, were proxied by the rs12916 variant; Evolocumab, targeting the *PCSK9* gene, was proxied by rs11591147; Ezetemibe, targeting the *NPC1L1* gene, was proxied by rs2073547^61^. Apo(a) inhibitors, targeting the *LPA* gene, were proxied by rs10455872^62^. *APOC3* inhibitors, targeting the *APOC3* gene, were proxied by rs10790162^61^. The effect of statins, Evolocumab and Ezetemibe on disease risk for CHD and type 2 diabetes were obtained from randomized controlled trials (RCTs)^20,21,28,29^ for comparison with the MR estimates, equivalent RCT results for *APOC3*-and apo(a)-inhibitors were not available (Supplementary table 4).

## Acknowledgements

We gratefully acknowledge all the studies and databases that have made GWAS summary data available (the investigators of these studies and databases did not participate in the analysis, writing or interpretation of this report): **ADIPOGen** (Adiponectin genetics consortium), **AMDGene** (Age-related Macular Degeneration Gene Consortium), **BioBank Japan Project**, **C4D** (Coronary Artery Disease Genetics Consortium), **CARDIOGRAM** (Coronary ARtery Disease Genome wide Replication and Meta-analysis), **CKDGen** (Chronic Kidney Disease Genetics consortium), **CORNET** (The CORtisol NETwork), **dbGAP** (database of Genotypes and Phenotypes), **DIAGRAM** (DIAbetes Genetics Replication And Meta-analysis), **ENIGMA** (Enhancing Neuro Imaging Genetics through Meta Analysis), **EAGLE** (EArly Genetics & Lifecourse Epidemiology Eczema Consortium, excluding 23andMe), **EGG** (Early Growth Genetics Consortium), **GABRIEL** (A Multidisciplinary Study to Identify the Genetic and Environmental Causes of Asthma in the European Community), **GCAN** (Genetic Consortium for Anorexia Nervosa), **GEFOS** (GEnetic Factors for OSteoporosis Consortium), **GIANT** (Genetic Investigation of ANthropometric Traits), **GIS** (Genetics of Iron Status consortium), **GLGC** (Global Lipids Genetics Consortium), **GliomaScan** (cohort-based genome-wide association study of glioma), **GoT2D** and **T2D-GENES** consoritum, **GPC** (Genetics of Personality Consortium), **GUGC** (Global Urate and Gout consortium), **HaemGen** (haemotological and platelet traits genetics consortium), **HRgene** (Heart Rate consortium), **IAC** (the International Aneurysm Consortium), **ICBP** (International Consortium for Blood Pressure), **IGAP** (International Genomics of Alzheimer’s Project), **IIBDGC** (International Inflammatory Bowel Disease Genetics Consortium), **ILCCO** (International Lung Cancer Consortium), **ImmunoBase** (resource focused on the genetics and genomics of immunologically related human diseases), **IMSGC** (International Multiple Sclerosis Genetic Consortium), **MAGIC** (Meta-Analyses of Glucose and Insulin-related traits Consortium), **MESA** (Multi-Ethnic Study of Atherosclerosis), **NHGRI-EBI GWAS catalog** (National Human Genome Research Institute and European Bioinformatics Institute Catalog of published genome-wide association studies), **PanScan** (Pancreatic Cancer Cohort Consortium), **PGC** (Psychiatric Genomics Consortium), **Project MînE** consortium, **ReproGen** (Reproductive Genetics Consortium), **SSGAC** (Social Science Genetics Association Consortium) and **TAG** (Tobacco and Genetics Consortium), **TRICL** (Transdisciplinary Research in Cancer of the Lung consortium).

We gratefully acknowledge the assistance of Dr Johannes Kettunen.

## Funding

Supported by Cancer Research UK grant C18281/A19169 (the Integrative Cancer Epidemiology Programme) and the Roy Castle Lung Cancer Foundation (2013/18/Relton). The Medical Research Council Integrative Epidemiology Unit is supported by grants MC_UU_12013/1, MC_UU_12013/2 and MC_UU_12013/8. PCH is supported by a Cancer Research UK Population Research Postdoctoral Fellowship (C52724/A20138). Jack Bowden is supported by a MRC Methodology Research Fellowship (grant MR/N501906/1). D.M.E is supported by an Australian Research Council Future Fellowship (FT130101709).

